# Hypothalamic Vasopressin Neurons Enable Maternal Thermoregulatory Behaviors

**DOI:** 10.1101/2025.01.23.634569

**Authors:** Zahra Adahman, Rumi Ooyama, Dinore B. Gashi, Zeyneb Z. Medik, Hannah K. Hollosi, Biswaranjan Sahoo, Nana D. Akowuah, Justin S. Riceberg, Ioana Carcea

## Abstract

Newborns of many mammalian species are partial poikilotherms and require adult thermoregulatory care for survival. In mice, pup survival in cold and cool ambient temperature depends on the ability of adult caregivers to huddle pups and bring them into a high-quality nest. It is therefore essential that adult mice adjust parental care as a function of changes in ambient temperature. Here, we investigated how mouse maternal care adapts to a range of temperatures, from cold to warm. We show that changes in ambient temperature affect several individual and co-parenting maternal behaviors in both dams and virgin female mice, and modulate activity of vasopressin neurons. Furthermore, we establish that the effects of ambient temperature on both maternal care and the activity of vasopressin neurons depend in part on thermosensation, specifically on the TRPM8 sensor. Using trans-synaptic anterograde tracing and whole-brain activity mapping, we find that vasopressin neurons from the paraventricular hypothalamic nucleus connect synaptically with temperature-responsive brain structures implicated in maternal care. We then show that optogenetic activation of vasopressin projections to the central amygdala, a structure activated by cold ambient temperature, recapitulates the effects of cold on co-parenting behaviors. Our data provide a biological mechanism for maternal thermoregulatory behavior in mice with translational relevance to the reported association between ecosystem temperature fluctuations and variations in human child neglect cases.

## INTRODUCTION

Changes in ambient temperature associate with the incidence of child maltreatment in humans, with fewer cases of child neglect in cold and more cases in warm temperature [1]. This link could emerge as a consequence of thermoregulation. Endothermic animals developed complex physiological and behavioral adaptations to regulate body temperature [2–5]. In many species additional behaviors evolved in adults to maintain constant core body temperature of poikilothermic infants [6]. In mice, these thermoregulatory behaviors are an essential part of the maternal care repertoire, and include nest building, retrieving pups into the nest, and huddling with pups in nest [7, 8]. Previous work, primarily in rat dams, has shown that cold ambient temperature stimulates pup interactions and nest building, whereas warm temperature decreases time spent in nest [6, 9, 10]. However, the mechanisms underlying these adaptations remain unknown.

Sensing ambient temperature begins with temperature receptors expressed on sensory fibers or on keratinocytes. The majority of temperature receptors are cation channels in the <u>T</u>ransient <u>R</u>eceptor <u>P</u>otential (TRP) family that are gated by ambient temperature. Several TRP channels respond to changes in temperature within the non-noxious temperature range (16°-38°C). TRPM8 opens at temperatures below 25°C and is necessary for cold and cool sensing [11, 12]. However, TRPM8 also plays a role in warm processing, as mice lacking TRPM8 (TRPM8-/-) have deficits in perception of both cold and warm temperatures [13–15]. Information about temperature travels from peripheral sensory neurons to the central nervous system reaching the thalamo-cortical pathway for temperature perception, and the ponto-hypothalamic circuit for coordinated thermoregulatory physiology and behavior [2, 3, 16, 17]. Temperature responsive neurons in the ponto-hypothalamic pathway project to several structures that are important for maternal behavior, including the paraventricular nucleus of the hypothalamus (PVN) [18, 19]. Prior research on maternal care has highlighted PVN as a key brain structure, mediating nest building, pup retrieval, and time spent in nest [20–22]. Within PVN, several peptides have been shown to play important roles in maternal behavior, primarily oxytocin (OT) and vasopressin (AVP) [8]. Whereas the role of oxytocin neurons in enabling maternal behavior in dams and virgin females has been extensively studied [20, 21, 23–25], the role of AVP+ neurons is less well understood. PVN-AVP+ neurons promote pup retrieval and huddling in nest [26], potentially by synapsing on *Galanin*-expressing neurons in the medial preoptic area (MPO), another major regulator of maternal care [7, 27–30]. AVP has also been linked to nest building behavior [31], and to intruder-directed maternal aggression by acting on the central amygdala (CEA) [32].

Here, we characterize how acute (maximum 4 hours) exposure to a range of non-noxious ambient temperatures modulates different maternal behaviors in primiparous and nulliparous female mice. We show that thermoreceptors implicated in self-directed thermoregulation also serve maternal thermoregulatory behaviors directed towards pups. Finally, we find that AVP+ neurons in PVN respond to ambient temperature and mediate co-parenting but not individual maternal thermoregulatory behaviors via projections to the CEA. Taken together, our study identifies biological mechanisms by which mouse maternal care adapts to changes in ambient temperature, and provides insights into how ecosystem fluctuations influence maternal care in mammals.

## RESULTS

### Dams and virgin surrogates exhibit maternal thermoregulatory behaviors

To investigate how maternal behaviors of dams and surrogate virgin females adapt to changes in ambient temperature, we employed a sequence of previously described behavioral tests: 1. Four hours of co-housing between dam, virgin and P1-P5 pups, with access to nest, food and water. 2. Pup retrieval testing of individual adults (with two P5-P7 pups). 3. Nest building assay for individual adults in the presence of three pups (P7-P14). All behavioral tests were video recorded for behavioral analysis. The initial co-housing also served as priming for virgins, as they gained experience with caring for pups and became maternal, as we previously described [23]. We tested these behaviors at different non-noxious ambient temperatures: cold (CT, 15-17°C), room temperature (RT, 20-22°C), thermoneutral temperature (TN, 29-31°C) and warm temperature (WT, 36-38°C). We used 70 dams and 79 female virgins who performed at least one of the behavioral tests only once, at one of the four temperatures considered. For some tests we had to exclude data that was identified as outlier, did not meet criteria, or where data collection was compromised (see **Methods**).

In the co-housing task we characterized both individual, and dam-virgin co-parenting behaviors. We quantified the time each animal spends in the nest with pups (**Figure 1A**). We find a significant effect of temperature in dams (**Figure 1Ai**, one-way ANOVA, F(3,25) = 13.44, p < 0.0001, N = 29) and in virgins (**Figure 1Aii**, F(3,25) = 4.36, p = 0.013, N = 29). Pairwise comparison with the RT condition (habitual temperature in the mouse vivarium) shows that dams spend significantly less time in nest when co-housing is done at WT (Dunnett’s multiple comparison test, p = 0.0002), and virgins spend significantly more time in nest during CT (p = 0.017). Time in nest for all dams and virgins was inversely correlated with ambient temperature (**Figure 1Aiii**, Pearson correlation, r = −0.56, p < 0.0001). We also find that ambient temperature modulated co-nesting, where dam and virgin spend more time together in nest in CT and less time in WT (**Figure 1Aiv**, F(3,24) = 15.61, p < 0.0001; RT vs CT, p = 0.005; RT vs WT, p = 0.016).

**Figure 1:**
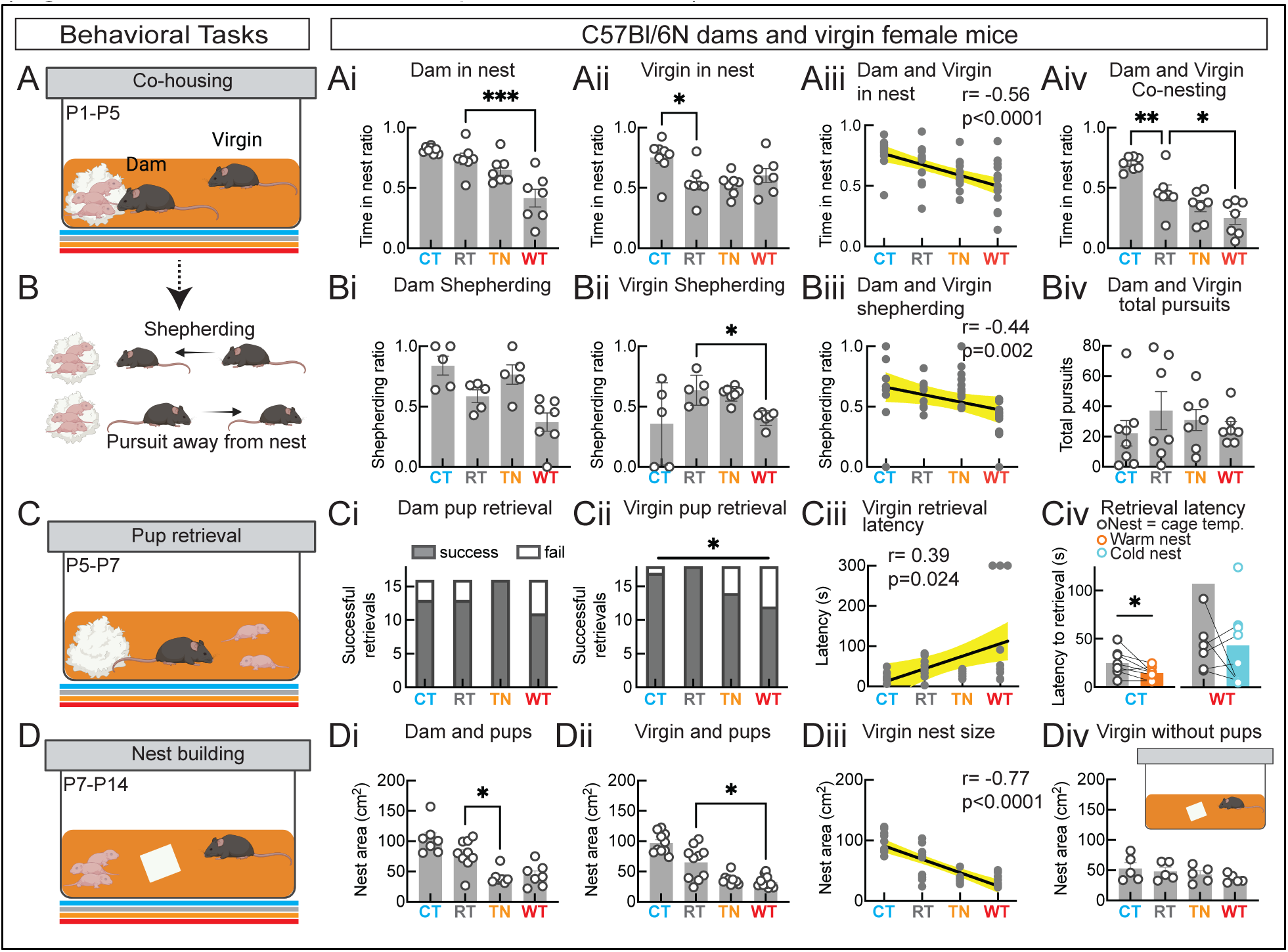
Maternal thermoregulatory behaviors in C57Bl/6N females. **A**. Schematic of the cohousing behavior task. **Ai**. The ratio of time spent in nest by dams. **Aii**. The ratio of time spent in nest by adult virgin females. **Aiii**. Correlation between the ratio of time spent in nest and ambient temperature. **Aiv**. The ratio of time spent co-nesting between dams and virgins at different ambient temperatures. **B**. Depiction of shepherding and pursuits away from nest. **Bi**. The ratio of shepherding by dams. **Bii**. The ratio of shepherding by virgins. **Biii**. Correlation between the ratio of shepherding and ambient temperature. **Biv**. Frequency of all pursuits (toward and away from nest). **C**. Schematic of the pup retrieval behavior task. **Ci.** Dam pup retrieval success. **Cii.** Virgin pup retrieval success. **Ciii.** Correlation of pup retrieval latency in virgins and ambient temperature. **Civ.** Virgin pup retrieval in ‘relief’ nest. **D.** Schematic of the nest building behavioral assay. **Di**. Area of nests built by dams. **Dii**. Area of nests built by virgins. **Diii**. Correlation of nest area and ambient temperature. **Div.** Area of nests built by virgins in the absence of pups. *p < 0.05, **p < 0.01, ***p < 0.005. Shaded areas in Aiii, Biii, Ciii, and Diii represent 95% confidence intervals.

We previously described an interaction between dam and virgin during the co-housing procedure which we called shepherding, where dams pursuit the virgin towards the nest with pups (**Figure 1B**, [23]). Here we show that the ratio of dam shepherding (out of all pursuits) is sensitive to changes in ambient temperature (**Figure 1Bi**, one-way ANOVA, F(3,18) = 8.54, p = 0.001, N = 22). Furthermore, we show that virgins can also shepherd the dams, and the ratio of virgin shepherding is sensitive to changes in ambient temperature (**Figure 1Bii**, Welch’s ANOVA, W(3,8.83) = 11.72, p = 0.002, N = 23). Pairwise comparison shows that WT significantly decreases shepherding behavior in virgins (Dunnett’s T3 test, RT vs WT, p = 0.024), indicating that pursuits away from nest predominate at WT. Overall, the ratio of shepherding for dams and virgins is inversely correlated with ambient temperature (**Figure 1Biii**, Spearman correlation, r = −0.44, p = 0.002). Ambient temperature does not have a significant effect on pursuit activity, as the number of total pursuits during co-housing did not change with changes in ambient temperature (**Figure 1Biv**, Kruskal-Wallis test, p = 0.689, N = 29 pairs).

We next investigated pup retrieval behavior. Despite a trend for WT to decrease performance, pup retrieval success was not significantly affected by temperature in dams (Fisher’s exact test, p=0.115). In virgins, we found a significant effect of temperature on success of pup retrieval behavior, with fewer retrievals in WT (p = 0.017). The latency with which virgins retrieved the first pup linearly correlates with ambient temperature, indicating that the colder the environment the faster virgins retrieve (**Figure 1Ciii**, Spearman correlation, r = 0.39, p = 0.024). To determine if virgins bring pups into the nest to thermoregulate the pups or themselves, we repeated the experiment at CT and WT while providing a ‘relief’ nest: thermoneutral nest in CT environment, and cool nest in WT environment. We find that at CT, virgins retrieve pups faster in the relief nest compared to the nest of same temperature as the environment (**Figure 1Civ**, paired t-test, t(df) = 2.54 (7), p = 0.038, N = 8 pairs). On the contrary, the presence of the relief nest at WT did not significantly speed up pup retrieval (**Figure 1Civ**, Wilcoxon matched-pairs signed rank test, W = −20, p = 0.195, N = 8 pairs).

Lastly, we investigated how changes in ambient temperature affected nest building behavior in the presence of pups (**Figure 1D**). The area of the nest built for pups depended on ambient temperature in both dams (**Figure 1Di**, Kruskal-Wallis test, p = 0.0004, N = 30) and virgins (p < 0.0001, N = 40). Compared to the RT condition, dams built smaller nests at TN (Dunn’s test, p = 0.044), and virgins built smaller nests at WT (p = 0.043). The area of the nest built by virgins inversely correlates with ambient temperature (Spearman correlation, r = −0.77, p < 0.0001). Similarly, the nest quality depended on ambient temperature, as both dams and virgins built lower score nests at higher temperatures (**Figure S1**). Importantly, temperature does not have a significant effect on nest size or quality in the absence of pups, indicating that the effort to build larger and better nests at cold and cool temperatures is motivated by the presence of pups (**Figure 1Div**, Kruskal-Wallis test, p = 0.227, N = 20).

Taken together, the results from the above experiments indicate that female mice adapt pup-directed behaviors to provide maternal thermoregulatory care. The ethological aspects of maternal thermoregulation that we tested do not depend on biological changes associated with gestation, parturition and lactation, as we found similar effects of ambient temperature changes in dams and virgin females. Therefore, to identify biological substrates for maternal thermoregulatory behaviors, we next investigated the contribution of thermosensory mechanisms in female virgin surrogates.

### TRPM8 -/- mice have impaired maternal thermoregulation

Several mechanistic models could account for the maternal behavior adaptations to ambient temperature. To test if peripheral sensing of ambient temperature plays a role in initiating maternal thermoregulatory behavior, we used mice lacking TRPM8, a cold receptor expressed primarily by sensory fibers [11, 12]. TRPM8 is expressed by both cold-activated and warm-suppressed sensory fibers (**Figure 2A**, [14]) and prior work has shown that TRPM8-/- mice have deficits in both cold and warm sensing [13–15], making this mouse line ideal for investigating the effects of temperature on maternal behavior (**Figure 2A**).

**Figure 2:**
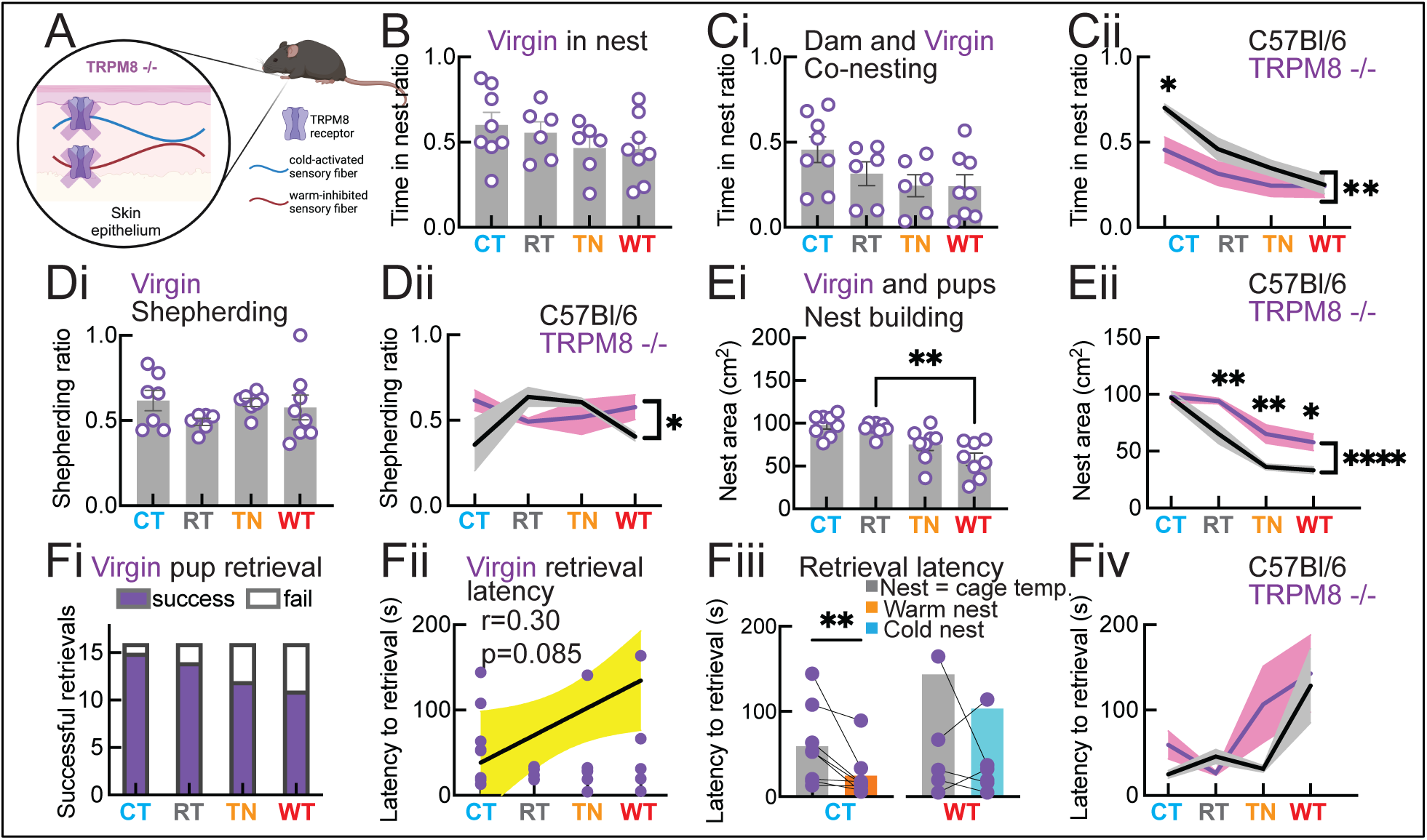
Maternal behavior in TRPM8 -/- virgin females. **A**. Schematic of TRPM8 expression on both cold-activated and warm-suppressed sensory fibers. **B**. The ratio of time spent in nest by TRPM8 -/- virgins during cohousing with C57Bl/6N dams and pups. **Ci**. The ratio of time spent co-nesting. **Cii**. Time co-nesting by TRPM8 -/- vs C57Bl/6N virgins. **Di**. Ratio of shepherding. **Dii**. Shepherding ratio by TRPM8 -/- vs C57Bl/6N virgins. **Ei**. Nest area. **Eii**. Nest area built by TRPM8 -/- vs C57Bl/6N virgins. **Fi.** Pup retrieval success. **Fii.** Correlation between retrieval latency and ambient temperature. **Fiii.** Pup retrieval latency when relief nest is provided. **Fiv.** Latency of pup retrieval by TRPM8 -/- vs C57Bl/6N virgins. *p < 0.05, **p < 0.01, ****p < 0.001. Shaded area in Fii represents 95% confidence intervals.

In the co-housing experiment, we used TRPM8 -/- virgin females and C57Bl/6N dams and pups. We found that ambient temperature had no significant effect on the time TRPM8 -/- females spent in nest (**Figure 2B**, one-way ANOVA, F(3,24) = 1.06, p = 0.381, N = 28). Similarly, temperature did not affect the time TRPM8 -/- virgins spent co-nesting with C57Bl/6N dams and pups (**Figure 2Ci**, F(3,24) = 2.18, p = 0.115, N = 28). Co-nesting behavior differs between C57Bl/6N and TRPM8 -/- virgins (**Figure 2Cii**, two-way ANOVA, effect of genotype, F(3,48) = 8.40, p = 0.005, N = 56). Pairwise comparisons showed that TRPM8 -/- virgins spend less time co-nesting with dams and pups at CT (Benjamini-Krieger-Yekutieli multiple comparison test for C57Bl/6N vs TRPM8 -/- co-nesting, p = 0.005 at CT, p = 0.113 at RT, p = 0.255 at TN, and p = 0.909 at WT). Shepherding ratio in TRPM8 -/- mice was not affected by temperature (**Figure 2Di**, one-way ANOVA, F(3, 24) = 1.007, p = 0.406, N = 28), leading to a different pattern compared to C57Bl/6N virgin females (**Figure 2Dii**, two-way ANOVA, genotype X temperature interaction, p = 0.021, N = 50).

Temperature changes still affected nest building behavior in TRPM8 -/- virgins, as they built smaller nests at WT compared to RT (**Figure 2Ei**, F(3,28) = 10.66, p < 0.0001, N = 32; Dunnett’s test for multiple pairwise comparison, RT vs WT, p = 0.0003). However, nest building differs between C57Bl/6N and TRPM8 -/- virgin females (**Figure 2Eii**, effect of genotype in two-way ANOVA is F(1,64) = 25.65, p < 0.0001, N = 72). In pairwise comparisons we find that TRPM8 -/- mice build larger nests than C57Bl/6N mice at all temperatures except CT (RT, p = 0.003; TN, p = 0.003; WT, p = 0.016). We found similar effects on nest quality (**Figure S1**).

Temperature does not affect pup retrieval success in TRPM8 -/- mice (**Figure 2Fi**, Fisher’s exact test, p = 0.315). Similarly, retrieval latency does not linearly correlate with ambient temperature (**Figure 2Fii**, Spearman correlation, r = 0.23, p = 0.218, N = 30). However, we find that the presence of a ‘relief’ nest in the CT but not WT condition speeds up retrieval in TRPM8 - /- virgins similarly as in C57Bl/6 virgins (**Figure 2Fiii**, Wilcoxon test, CT, p = 0.007; WT, p =0.562). Moreover, a genotype comparison shows no difference in retrieval latency between TRPM8 -/- and C57Bl/6N virgins, as both show similar patterns of temperature modulation (**Figure 2Fiv**, two-way ANOVA main effect of temperature F(3,55) = 4.87, p = 0.004, N = 63).

Thus, TRPM8 -/- mice have deficits in maternal behavior adaptation to both cold temperatures (time in nest, shepherding), as well as thermoneutral and warm temperatures (nest building). TRPM8 -/- mice have minimal disruptions of temperature-dependent pup retrieval, indicating that this behavior could rely on additional mechanisms other than temperature detection. Indeed, we confirmed prior findings showing that cold modulates pup vocalizations (**Figure S2**), vocally communicated triggers of maternal pup retrieval behavior [20, 33, 34]. Our findings validate a role for thermosensation in specific aspects of maternal thermoregulatory behavior.

### Ambient temperature modulates activity of hypothalamic AVP+ neurons

Temperature information travels from peripheral sensory fibers to spino-ponto-hypothalamic circuits that mediate physiological and behavioral self-thermoregulatory responses [3]. PVN is one of the hypothalamic nuclei important for thermoregulation [35–37]. PVN also hosts OT and AVP expressing neurons, that have been linked to maternal behavior in both dams and virgin female mice [20–22, 26, 31, 32]. To determine if ambient temperature modulates activity of OT+ and AVP+ neurons in PVN, we investigated the expression of c-fos immediate early gene. Virgin females were exposed to one of the four temperature conditions for four hours, and then the brains were collected for c-fos and peptide immunostaining (**Figure 3A**). We find an effect of temperature on c-fos expression in PVN-OT+ neurons that was driven by suppressed expression at TN (**Figure 3B**, one-way ANOVA F(3,12) = 5.46, p = 0.013, N = 16; Dunnett’s test for RT vs CT, p = 0.999; RT vs TN, p = 0.041; RT vs WT, p = 0.582).

**Figure 3:**
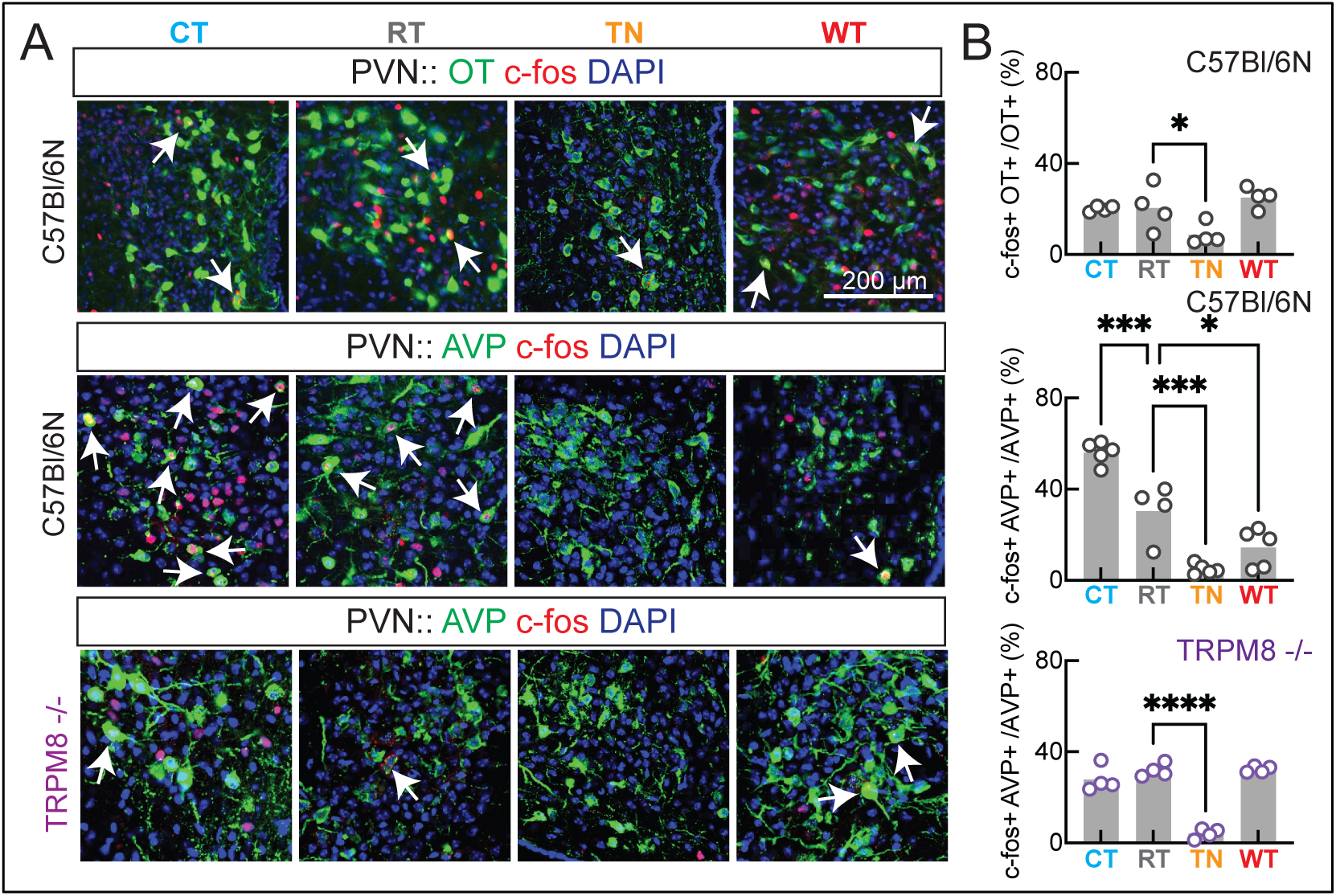
Temperature modulation of c-fos expression in PVN neuroendocrine cells in C57Bl/6N and TRPM8 -/- virgin female mice. **A.** Representative images showing c-fos, OT+ and AVP+ after incubation at different temperatures. **B.** Quantification of c-fos expression in neuroendocrine cells. *p < 0.05, ***p < 0.001, ****p < 0.0001.

PVN-AVP+ neurons showed more profound responses to ambient temperature, as they were activated at CT and suppressed at TN and WT (F(3,15) = 43.52, p < 0.0001, N = 19; RT vs CT, p = 0.0004, RT vs TN, p = 0.0004, RT vs WT, p = 0.016). Importantly, both the activation of PVN-AVP+ neurons at CT and their suppression at WT were absent in TRPM8 -/- female mice (F(3,12) = 60.65, p < 0.0001, N = 16; RT vs CT, p = 0.291; RT vs TN, p < 0.0001, RT vs WT, p = 0.994). These findings indicate that PVN-AVP+ neurons could mediate thermosensation-driven maternal thermoregulatory behaviors.

### Temperature responsive structures downstream of PVN-AVP+ cells

To investigate if PVN-AVP+ neurons could play a role in maternal thermoregulatory behaviors, we started by investigating whether they communicate synaptically with structures that play a role in maternal care and respond to changes in ambient temperature. We studied this in two steps. First, we mapped anterograde trans-synaptic connections from PVN-AVP+ neurons. For this, we injected in the PVN of AVP-IRES-Cre mice a modified herpes virus that transduces TdTomato fluorophore in Cre-expressing cells and then crosses the synapses to post-synaptic neurons. After 48 hrs of incubation we identified labeled cells in several structures that contribute to maternal behavior: medial preoptic area (MPO, **Figure 4Ai**), ventromedial hypothalamus (VMH, **Figure 4ii**), ventral tegmental area, dorsal raphe, and periaqueductal gray (**Table 1**).

**Figure 4:**
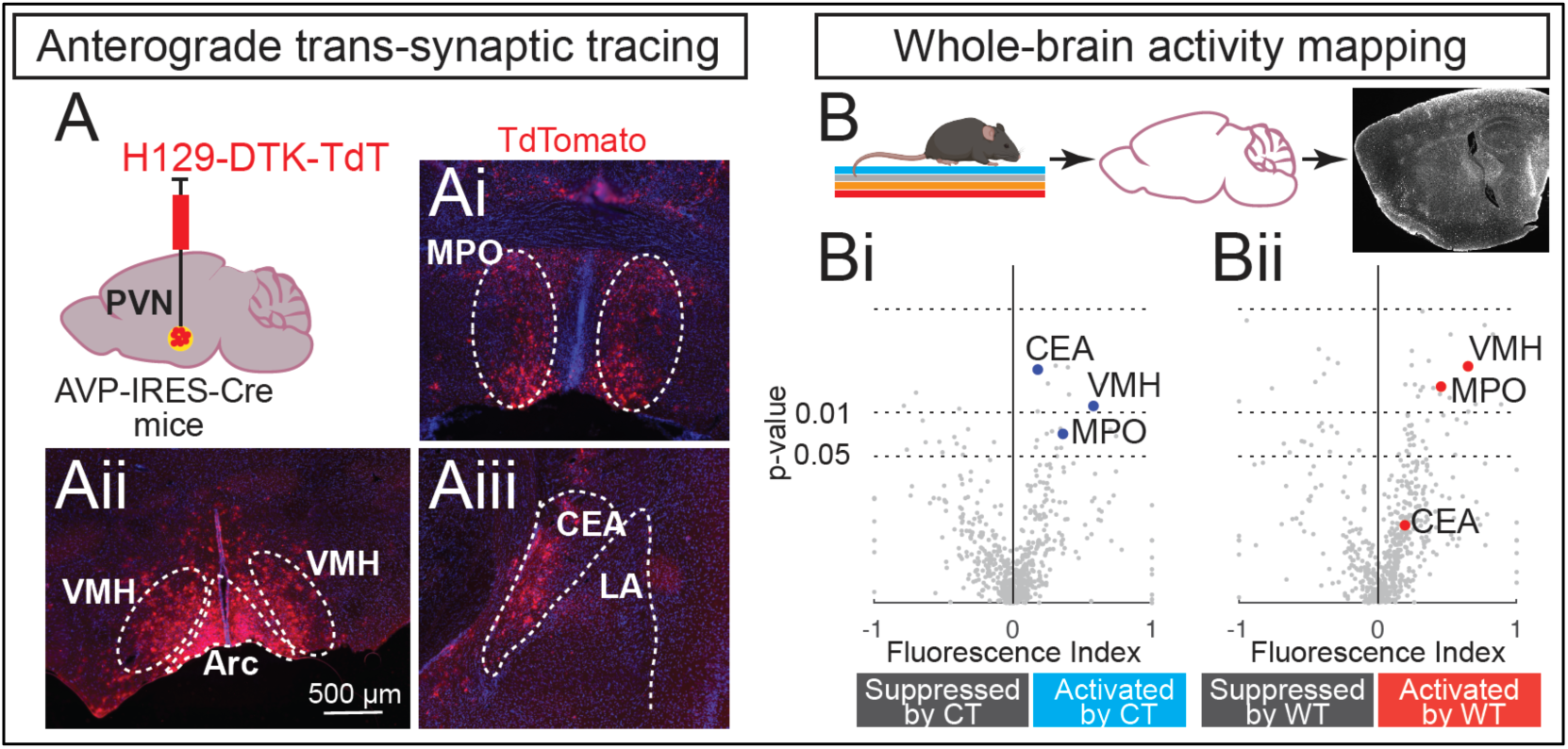
Temperature modulation of immediate early gene expression in brain regions receiving PVN-AVP synaptic input. **A-Aiii.** Experimental diagram and representative images showing anterograde trans-synaptic tracing from PVN-AVP+ neurons. **B-Bii.** Quantification of whole-brain activity mapping for Npas4 immediate early gene.

**Table 1:** Quantification of anterograde trans-synaptic tracing from PVN-AVP+ neurons (unilateral injections). + = 0.1-100 cells in 1 x 10^6^ px^2^; ++ = 100.1-200 cells in 1 x 10^6^ px^2^; +++ = 200.1-400 cells in 1 x 10^6^ px^2^; ++++ = 400.1-above cells in 1 x 10^6^ px^2^.

| No. | Regions with TdTomato+ cells | Abbreviation | Laterality | Intensity |
| --- | --- | --- | --- | --- |
| 1 | dorsal transition zone | DTr | Bilateral | + |
| 2 | ventral orbital cortex | VO | Bilateral | + |
| 3 | medial orbital cortex | MO | Bilateral | + |
| 4 | cingulate cortex, area 2 | Cg2 | Bilateral | +++ |
| 5 | nucleus accumbens core and shell | Nac | Bilateral | + |
| 6 | anteroventral periventricular nucleus | AVPe | Bilateral | ++ |
| 7 | lateral septal nucleus, ventral part | LSV | Bilateral | ++ |
| 8 | mediodorsal thalamus | MD | Unilateral | ++++ |
| 9 | paraventricular thalamic nucleus | PVA | Bilateral | +++ |
| 10 | suprachiasmatic nucleus | SCh | Bilateral | ++ |
| 11 | paraventricular nucleus of the hypothalamus | PVN | Bilateral | +++ injection site |
| 12 | nucleus of the stria medularis | sm | Unilateral | ++++ |
| 13 | anterior hypothalamic area | AH | Bilateral | + |
| 14 | supraoptic nucleus | SO | Bilateral | ++ |
| 15 | median preoptic area | MnPO | Bilateral | ++ |
| 16 | bed nucleus of stria terminalis | STS | Bilateral | ++++ |
| 17 | lateral hypothalamic nucleus | LH | Bilateral | ++ |
| 18 | anterodorsal thalamic nucleus | AD | Unilateral | ++++ |
| 19 | medial preoptic area | MPA | Bilateral | + |
| 20 | CA2 of hippocampus | CA2 | Bilateral | + |
| 21 | central amygdala | CeA | Bilateral | ++ |
| 22 | arcuate nucleus | Arc | Bilateral | +++ |
| 23 | retrochiasmatic area | RCh | Bilateral | ++ |
| 24 | median eminence | ME | Bilateral | + |
| 25 | lateral amygdala | La | Bilateral | + |
| 26 | enthorinal cortex | Ent | Bilateral | + |
| 27 | ectorhinal cortex | Ect | Bilateral | + |
| 28 | primary auditory cortex | Au1 | Bilateral | + |
| 29 | secondary visual cortex | V2 | Bilateral | ++ |
| 30 | lateral mammillary nucleus | LM | Bilateral | ++ |
| 31 | CA3 of posterior hippocampus | CA3 | Bilateral | ++ |
| 32 | peri-aqueductal gray | PAG | Bilateral | ++ |
| 33 | dorsomedial hypothalamic nucleus | DMH | Bilateral | + |
| 34 | posterior hypothalamus | PH | Bilateral | + |
| 35 | ventral tegmental area | VTA | Bilateral | ++ |
| 36 | substantia nigra pars compacta | SNC | Bilateral | + |
| 37 | dorsal raphe nucleus | DR | Bilateral | ++++ |
| 38 | A5 noradrenaline cells | 5ADr | Unilateral | + |
| 39 | interfascicular nucleus | IF | Bilateral | + |
| 40 | retromammillary nucleus | RM | Bilateral | + |
| 41 | subgeniculate nucleus of prethalamus | SubG | Bilateral | + |
| 42 | lateral parabrachial nucleus | LPB | Bilateral | ++++ |

Second, we performed whole brain immediate early gene activity mapping following exposure to different temperatures. Virgin females were incubated alone at one of the four different temperatures for one hour, and after collection their brain was cleared, stained for Npas4 immediate early gene and imaged with light-sheet microscopy (**Figure 4B**). We calculated a fluorescence index to identify brain regions that had changes in Npas4 expression at CT and WT compared to RT condition ((F_CT_ – F_RT_)/(F_CT_ + F_RT_) and (F_WT_ – F_RT_)/(F_WT_+F_RT_)). Several brain regions were activated or suppressed by exposure to CT or WT (**Figure 4Bi, Bii,** uncorrected one sample t-test, H0 = 0). Few of these regions that receive PVN-AVP synaptic input (**Figure 4A**) were activated by changes in temperature: MPO and VMH were activated at both CT and WT, whereas CEA was activated only at CT. These findings indicate that PVN-AVP+ neurons project to brain structures that respond to changes in ambient temperature, and indicate potential mechanisms supporting maternal thermoregulatory behavior. Given our findings that only CT activates PVN-AVP+ cells, we next focused on PVN-AVP+ projections to CEA, which was specifically activated at CT and which previously received limited attention in the context of maternal behavior.

### PVN-AVP+ fibers in central amygdala support maternal thermoregulatory care

To determine if vasopressin innervation of CEA could contribute to maternal thermoregulation, we optogenetically activated PVN-AVP+ fibers in the CEA and tested maternal behavior at RT (**Figure 5A,B**). In AVP-IRES-Cre virgin females transducing Cre-dependent channelrhodopsin-2 E123T (ChETA) in PVN, we found increased time co-nesting and enhanced shepherding behavior after receiving blue light stimulation in CEA (hϖ ON) compared to the control condition (hϖ OFF) (**Figure 5C**i: paired t-test t(df) = 3.86, 4), p = 0.018, N = 5; **Figure 5Di**: t(df) = 3.89 (4), p = 0.017, N = 5). These behaviors were not changed by blue light stimulation in animals transducing a control virus (**Figure 5Cii**: t(df) = 2.45 (3), p = 0.091, N =4; **Figure 5Dii**: t(df) = 1.14 (3), p = 0.334, N = 4). Activation of PVN-AVP+ fibers in CEA only modulated co-parenting behaviors (co-nesting and shepherding), whereas individual maternal behaviors (overall time in nest, pup retrieval, and nest building) were not changed by stimulation (**Figure Ei**: t(df) = 2.27 (4), p = 0.085, N = 5; **Figure 5Fi**: t(df) = 0.75 (5), p = 0.486, N = 6; **Figure 5Gi**: Fisher’s exact test, p > 0.99), similar as in control animals (**Figure 5Eii**: t(df) = 2.18(3), p = 0.116, N = 4; **Figure 5Fii**: t(df) = 0.75(3), p = 0.507, N = 4; **Figure 5Gii**: p > 0.99).

**Figure 5:**
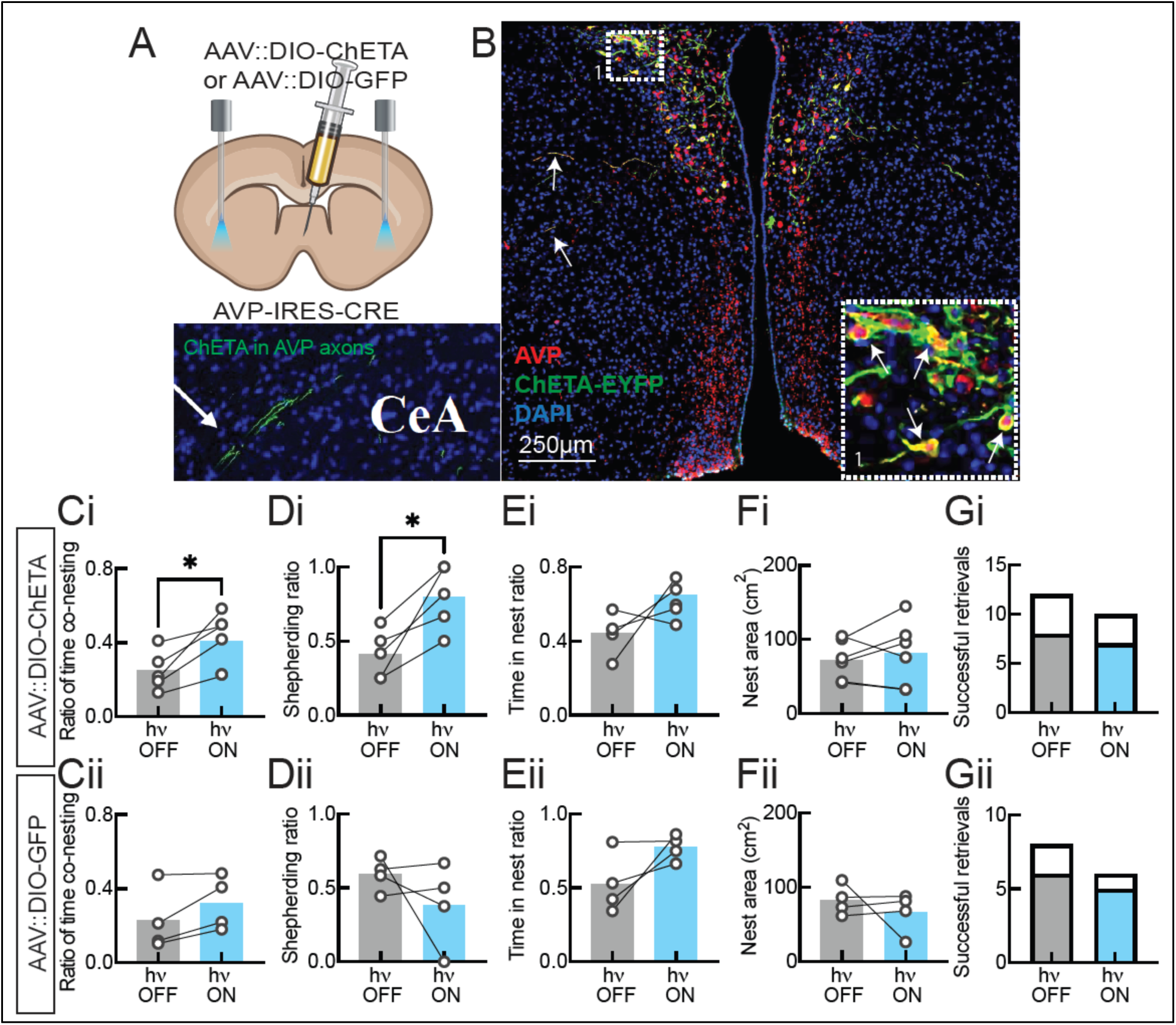
Optogenetic activation of PVN-AVP axons in CEA during behavior testing at RT. **A.** Schematic of viral injection and blue light stimulation. **B.** Example ChETA expression in AVP neurons in PVN and in fibers innervating CEA. **Ci,ii.** Co-nesting behavior. **Di,ii.** Shepherding behavior. **Ei,ii.** The ratio of overall time spent in nest by virgin. **Fi,ii.** Nest building behavior. **Gi,ii.** Successful pup retrievals. *p < 0.05.

## DISCUSSION

We found that maternal behavior adapts to changes in ambient temperature, with cold increasing and warm temperature decreasing several measures of ethological pup-directed care. Similarly, cold activates and warm suppresses hypothalamic PVN neurons producing AVP, a neurohormone implicated in maternal behavior. Both behavioral and hypothalamic AVP+ activity patterns were dependent on thermosensory mechanisms. We show that PVN-AVP+ neurons connect synaptically with brain structures responsive to changes in ambient temperature. Among these, CEA is specifically responsive to cold, and optogenetic stimulation of PVN-AVP+ fibers in CEA recapitulates the effects of cold temperature on co-parenting but not individual maternal thermoregulatory behaviors.

### Maternal thermoregulation

Generally, cold ambient temperature promoted behaviors that keep pups warm (nest building, huddling in the nest, pup retrieval) whereas warm temperature decreased these behaviors. We did not observe aggression behavior directed towards pups. Our results enrich the understanding of rodent maternal thermoregulatory behaviors [6, 9, 10], and mirror epidemiological findings in humans, where cold days associate with low and warm days with high numbers of child neglect cases [1]. Maternal thermoregulatory behaviors do not appear to be dependent on mating, gestational and lactation status, as both primiparous dams and nulliparous virgin surrogates display similar, albeit not identical, temperature-driven adaptations in pup care (**Figure 1**). Our data (not included in this manuscript) indicate that the estrus cycle does not significantly affect maternal behavior adaptations to ambient temperature. However this aspect needs to be studied further in light of recent findings that the ratio of gonadal hormones play an important role in shaping maternal behavior when self-directed needs are pressing [38]. An important question that we began addressing in this manuscript and are actively investigating further, is the extent to which the described changes in behavior are driven by pup needs versus self needs. In the pup retrieval experiment, if female mice were primarily motivated by maintaining their own thermal comfort, they would spend more time in the relief nest resulting in longer pup retrieval latencies. However, we observe the opposite effect, where the presence of the relief nest accelerates pup retrieval in cold (**Figure 1Civ**). Similarly, cold ambient temperature drove females to build larger nests only in the presence of pups and not when housed without pups (**Figure 1Div**). Together, these data indicate that cold temperature motivates maternal behavior in female mice.

The described temperature-dependent adaptations in maternal care could have profound consequences on offspring development and well-being [39–41]. In this context, it is important to emphasize that our manuscript investigates maternal behavioral adaptations in response to relatively acute changes in ambient temperature. Recent work showed that mouse thermoregulatory physiological processes adapt during long-term exposure (days, weeks) to extreme ambient temperature [42]. It remains to be determined if maternal care could undergo similar acclimations during prolonged exposure to cold or warm ambient temperature.

### Thermosensory mechanisms

Mechanistically, we found that maternal thermoregulatory behaviors depend in part on thermosensation, as they were disrupted in mice lacking the TRPM8 temperature receptor (**Figure 2**). This was particularly evident for co-parenting behaviors in the co-housing assay and for the nest building assay. Intriguingly, TRPM8-mediated thermosensation appeared to mediate the effects of cold in co-parenting assay and the effects of warm temperature on nest building. Prior work has shown that TRPM8 is expressed on both cold-excited and warm-suppressed sensory fibers [14], and our findings indicate that these distinct pathways may differentially modulate activity in brain structures supporting specific maternal behaviors.

Alternative mechanisms could partially account for maternal thermoregulatory behaviors. One such alternative is that maternal behavior adapts to communication signals from pups [43]. Mouse pups emit ultrasonic isolation calls that drive pup retrieval behaviors in dams and naïve virgin [20, 23, 33, 34]. We find that cold changes the rate and frequency of pup calls (**Figure S2**), confirming prior work on the effects of temperature on pup vocalizations [44, 45]. It is possible that ambient temperature changes additional types of pup communication signals, for example body odor. Prior work has shown that olfactory cues make an essential contribution to pup retrieval behavior [46, 47]. We found mixed effects of TRPM8 deletion on pup retrieval at different temperatures (**Figure 2Fi-iv**): only a trend for females to retrieve more and faster in cold, but a significant acceleration of pup retrieval when a relief nest is provided. Together, these data indicate that thermosensation is not the only mechanism supporting the effects of ambient temperature on pup retrieval behavior. Although the effects of pup communication signals on nest building are less well understood, it is possible that they also contribute to cold-motivated nest building, as TRPM8 -/- virgins made similar size nests as C57Bl/6 virgins when exposed to cold (**Figure 2Eii**).

Changes in local brain tissue temperature could represent an additional mechanism by which changes in ambient temperature shape maternal behavior. Compared to humans, mice have less tight control of their core body temperature [48, 49]. Specifically, when ambient temperature rises above thermoneutrality, core body temperature slowly increases [50]. Many temperature receptors are also expressed in internal organs, including the brain, and could induce changes in neuronal excitability [51]. TRPM8, however, is primarily expressed in sensory fibers, and has only very sparse expression in the brain [11, 52]. This justifies the use of global TRPM8 -/- mice. Nevertheless, to exclude the possibility that even such sparse brain expression of TRPM8 could be responsible for maternal thermoregulatory behaviors, future work should test mice in which TRPM8 is deleted specifically from sensory fibers.

### Vasopressin modulation of maternal behavior

In prior research on maternal care, the paraventricular nucleus of the hypothalamus (PVN) emerged as one of the brain structures mediating nest building, pup retrieval and time spent in nest [20–23]. Within PVN, several neuropeptides have been shown to play important roles in maternal behavior, primarily OT and AVP. Although both AVP and OT have been implicated in a variety of social behaviors, AVP has been generally considered to play more important roles in males than in females, while OT has been more often associated with maternal behavior in females [8, 53–58]. Intriguingly, we find that in females changes in ambient temperature modulate c-fos immediate early gene expression in AVP+ more than OT+ cells in the PVN (**Figure 3**). This corroborates prior work showing that cold increases AVP mRNA transcripts in rat PVN [59, 60]. OT- and AVP-producing neurons in other brain regions could also be engaged by changes in ambient temperature. In exploratory data not included in this manuscript, we do not find substantial temperature-induced changes in c-fos expression in OT+ neurons from supraoptic and suprachiasmatic nuclei, however such changes could be present in other OT-producing structures that we did not probe here: e.g. bed nucleus of stria terminalis, accessory and tuberal nucleus [61, 62].

While temperature-modulation of AVP+ cell activity could be linked to physiological functions of the AVP peptide (e.g.: control of water balance, vasoconstriction) [63], it also supports changes in maternal behavior. Increased brain AVP concentrations promote pup retrieval and increase the time spent caring for pups [26], potentially by synapsing on *Galanin* neurons in the medial preoptic area (MPO), another major regulator of maternal care [27, 28, 30]. AVP has also been linked to nest building behavior, as increased AVP expression was associated with decrease nest building [31]. Although this relationship is surprising given our finding that PVN-AVP+ activity increases in cold temperature, it could indicate that the effects of AVP on nest building depend on the source and target of AVP release. For example, mice selectively bred as good nest builders had fewer AVP+ cells in the suprachiasmatic nuclei but not PVN [64].

PVN-AVP+ neurons project to the posterior pituitary to release AVP in the blood stream for peripheral actions such as vasoconstriction, and project centrally to many brain areas implicated in parental behavior: MPO, VMH, bed nucleus of stria terminalis, periaqueductal gray, ventral tegmental area and others (**Figure 4**) [27, 28, 30, 65–70]. Some of these structures, such as MPO and VMH, respond to both increases and decreases in ambient temperature (**Figure 4)**, by receiving projections from temperature-responsive neurons within the ponto-hypothalamic pathway [5, 18, 71–74]. As such, these structures could contribute to maternal thermoregulatory behavior independent of AVP neuromodulation. Future work will directly test this possibility by manipulating AVP terminals in these brain structures. In our current study, we elected to focus on the CEA, a structure that receives PVN-AVP synaptic input and also responds to ambient cold but not warm temperature (**Figure 4**). CEA receives direct projections from PVN-AVP+ neurons (**Figure 5C**) [75] and also expresses vasopressin receptors 1a in both rodents and primates [76–78]. We optogenetically activated bilateral PVN-AVP→CEA fibers in animals housed at room temperature and tested maternal care. This manipulation mimicked the effects of cold on specific maternal thermoregulatory behaviors, namely co-parenting behaviors, but not on pup retrieval and nest building (**Figure 5**). Our findings indicate a role for CEA in maternal care, a role that has been minimally investigated in mice. A previous study in lactating rats showed that AVP concentration in CEA increases in response to interaction with an adult virgin female, but it led to increased aggression toward the virgin [32]. This finding is in apparent contradiction with our results, where PVN-AVP→CEA stimulation increases co-nesting and shepherding between virgin and dam (**Figure 5Ci, Di**). However, species-specific differences in co-parenting strategies could clarify the role of AVP in CEA. Whereas female mice typically nest communally, with dams and other females rearing pups together, rat females rarely nest communally and preferentially with co-reared sisters [79]. Thus, AVP in CEA could play a role in enhancing the species-specific co-parenting strategy.

## MATERIALS AND METHODS

### Animals

Wildtype timed pregnant and virgin (nulliparous) adult female mice (C57Bl6N) were obtained from Taconic Laboratories. TRPM8^-/-^ homozygous mice were generated in the laboratory of Dr. David Julius and obtained from The Jackson Laboratories (stock #008198). These mice have a mixed C57Bl/6N and J background. AVP-IRES2-Cre heterozygous mice were generated in the Allen Institute for Brain Science by Dr. Hongkui Zeng and obtained from The Jackson Laboratories (stock # 023530). Experiments were performed with primiparous dams at postnatal day 1-14 and virgin adult female mice of 8-12 weeks of age. Mice were housed at 22 ± 2°C, except when other temperature was indicated in the text. Animals were housed under a 12-h light/dark cycle (lights on at 8:00 a.m.), and had free access to food and water. Newborn mice were allowed to freely breastfeed from their respective dams. Dams are single housed with their litter of pups until weaning age of (21 days). Young virgin females are group-housed unless specifically stated in the test for behavioral testing purpose. The animal care and experimental procedures were carried out in accordance with the NIH guidelines and approved by the Rutgers State University of New Jersey Institutional Animal Care and Use Committee (IACUC).

### Statistical Analysis

The software used for data processing was Microsoft excel. The software used for all analysis was GraphPad Prism versions 9 and 10. For all datasets, we excluded outliers by using the ROUT method (with Q = 1%). That was followed by assessing the normal distribution of the sample Shapiro-Wilk test. Samples that had normal distribution were analyzed with parametric tests, and those that didn’t have normal distribution were analyzed with non-parametric tests.

### Temperature treatments

All the temperature conditions in this work were within the non-noxious temperature ranges. The four temperature conditions were achieved with artificial cooling or heating of the testing apparatus. Warm temperature (WT, 35-38°C) and thermoneutral (TN, 29-32°C) conditions were created by placing a heating pad (RIOGOO Pet Heating Pad with Auto Power, Amazon.com) under covered cages (NOLDUS phenotyper® 4500 cage with floor area of 45cm x 45cm (18“ x 18”), or the innocage®, (INNO VIVE) 14.7“L x 9.2”W x 5.5“H (maximum), (37.3 x 23.4 x 14.0 cm (maximum). The cold temperature (CT) condition (15-16°C) was created by setting a refrigerated chamber or insulated space to 15°C. The room temperature condition (22-25°C) was the ambient room temperature during the experiment. When the ambient room temperature (RT) in the observation room was below 21°C the heating pad was place under covered cages to achieve temperatures between 22-25°C. The temperature gradient conditions used in the pup retrieval assay were achieved by placing a cooling pack in WT condition and placing an activated hand warmer (HotHands) in the CT condition under the nest area to reach TN temperature (29-32°C).

### Behavior assays

#### Maternal behaviors

The sequence of behavior assays performed were co-housing assay, followed by pup retrieval and lastly nest building. The co-housing assay was performed first to prime the virgin females to acquire maternal behavior from interactions with dams and pups. Although all animals underwent co-housing for priming, only in a subset of animals we videotaped this procedure for behavioral analysis.

#### Co-housing assay

The pregnant females and naive virgins were kept separate cages until used for this behavior assay. After the pregnant female underwent parturition becoming dams, the assay was conduction between postnatal day one through seven (P1-P7). The naïve virgins were approximately eight to twelve (8-12) weeks old during the assay. Co-housing of a naive virgin (young female) with a dam and her litter was conducted for 4-6 hours per day for 3-4 consecutive days. Each pair of naive virgin and dam with her litter of 3 pups were exposed to a temperature conditions randomly assigned. The naive virgins’ fur was bleached and dye pink with semi-permanent dye and marked on their tails for identification. Animals were transferred from their home cage to the phenotyper cage (NOLDUS) for the assay and were returned to their home cage after completing the assay. The nesting materials from the mother’s home cages was also placed in the phenotyper cage. The floor of the phenotyper cage was covered with abundant bedding material. We added food pellets in the feeder and a water bottle. We first placed the dam and her postnatal day 1 (P1) litter in the cage. The assay was recorded using Color GigE cameras on the Ethovison software (NOLDUS). The behaviors were manually scored using BORIS version 7.7 through 8.4 [80]. Shepherding behavior (pursuing the other adult animal toward the nest) was identified as previously described [23], if the animal being chased entered the nest. Pursuits away from nest were all pursuits that did not end up with the chased animal in the nest. To be included in the calculation of the ‘Shepherding ratio’, the animals had to have a minimum 2 pursuit.

#### Pup retrieval assay

This behavior assay was performed in dams and young virgin female subjects adapted from Dr. Catherine Dulac’s work [81]. The young virgin females were housed individually for 18-24 hours prior to pup retrieval assay. Testing was performed with two (2) pups aged p3-7 in the home cage with the home nest material. A 30 min habituation period to the ambient temperature condition was performed before the testing commenced. The two C57BL/6N pups were removed from the nest and placed in different corners opposite the nest. Each trial had a ten (10) minutes limit, five minutes to retrieve each pup. For the ‘relief nest experiment, the conditions were as follow. At CT, the nest area was warmed to thermoneutrality (T°nest = 28.12 ± 2.31) by placing a handwarmer under the nest., and at WT the nest area was cooled (T°nest = 16.41 ± 1.38) using a cool pack under the nest. The test was recorded using logitech c-290 webcam. The behaviors were scored manually using BORIS version 7.7 through 8.4.

#### Maternal nest building assay

Maternal nest building was performed using dams and naïve virgins. Dams with their litter were exposed to all the temperature conditions randomly assigned for 4 hours per day for 3-4 consecutive days within the second week postpartum (P8-P14). Each dam with culled litter to 5 pups was placed in an innocage® (INNO VIVE) which was covered during the recording. In the cage adequate bedding, food, and hydrogel (a substitute for water) was provided. However, nest material was not provided. The recording began with the last minute of the acclimation stage. After ten (10) minutes of acclimation, a square cotton nestlet (Ancare®) was placed in a cage. The recording was ended after four hours. When maternal nest building was performed with naïve virgins, they are grouped housed with other virgins and each tested for maternal nest building with 5 pups. The nestlet was removed and flattened on a black surface and imaged along with a ruler to calibrate the pixels for calculating the area and perimeter. The nest structure was scored based on how broken the material was. The nest score of 0 for ‘Barely broken up’, 1 for ‘10-20% slightly broken up’, 2 for ‘20-50% partly broken up’, 3 for ‘50-80% partly broken up’ and 4 for ‘80-100% completely broken up’.

### Pup behaviors

#### Pup isolation ultrasonic vocalization recording

Pups aged P2-7 were each isolated from home cage with dam and rest of the litter to a beaker lined with bedding set at the four temperature conditions. The AVISOFT Avisoft-Bioacoustics CM16/CMPA microphone was placed around 3-4 inches above the base. The pup’s ultrasonic vocalizations (USVs) were recorded for 120 seconds. The pups were marked and returned to the home cage. This was repeated until all the pups were recorded for multiple litters of pups. The audio files were analyzed using the MUPET algorithm [82].

### Adult-only behavior

#### Virgin nest building assay in the absence of pups

Grouped housed naive C57Bl/6N virgins were separated and randomly selected to a nest building assay. They were each placed in a single housed testing apparatus (innocage® (INNO VIVE)) which was covered during the recording. In the cage adequate bedding, food, and hydrogel (a substitute for water) was provided. However, nest material was not provided. After ten minutes of acclimation the recording began with a square cotton nestlet (Ancare®) was placed in a cage. No pups were present. The recording ended after four (4) hours. The nestlet was removed and flattened on a black surface and imaged along with a ruler to calibrate the pixels for calculating the area and perimeter. The nest structure was scored and analyzed same as in the maternal nest building assay.

### Immediate early gene expression assessment

#### Temperature exposure assay

Grouped housed naive C57Bl/6N virgins were habituated to handling and single housing in a testing apparatus (innocage® (INNO VIVE)) for four (4) hours over four days. On the fifth day (test day), they were each randomly selected to the same test apparatus set at one of the four temperature conditions. The temperature exposure lasted for four hours. The mice were perfused, and brain tissue was collected within five-fifteen minutes from the end of the temperature exposure.

#### Immunohistology

Brain tissues were collected after transcardial perfusion with 1 X PBS, followed with 4% paraformaldehyde (PFA). Tissues were further preserved in a paraformaldehyde solution for 1–2 h at 4 °C, followed by cryopreservation of tissue in 15% and then 30% sucrose solution at 4 °C. The tissues were embedded in OCT solution and sectioned at 20 µm thickness with a cryostat then mounted on positively charged slides. Sections were blocked and permeabilized with 5% goat serum solution with 0.1% triton-X in 1x PBS for 1 h at room temperature. Primary antibodies were diluted in 1% goat serum and 0.01% triton-X in 1x PBS solution and then applied for 24hr - 48hr at 4 °C. We used a mouse anti-oxytocin antibody (1:100, a gift from H. Gainer at national Institutes for Health) [83], mouse anti-vasopressin antibody (a gift from H. Gainer at national Institutes for Health, 1:100) [83], rabbit anti-c-fos (Cell Signaling Technology, 1:500), a chicken anti-GFP (Aves, 1:500), and a rabbit anti-tDT (Takara, 1:500) antibody. The sections were washed in 2 to 3X in PBS solution. Then, the appropriate solutions of fluorophore-conjugated secondary antibodies were applied for 1.5 h at room temperature. All secondary antibodies were from Jackson ImmunoResearch and used at 1:1000 (Anti-Rabbit Goat Cy3, Anti-Mouse Goat Cy2, and Anti-Chicken Goat Alexa Fluor 488). The sections were washed in 2 to 3X in PBS solution again. After which, all sections were counterstained with DAPI (4’,6- diamidino-2-phenylindole). Slides sealed with Fluoromount-G™ Mounting Medium (Invitrogen) and a glass cover coverslip. Slides were imaged using a Nikon A1R+ HD Confocal Microscope and a Nikon Ti2 HCA Inverted Fluorescence Microscope at Rutgers New Jersey Medial School, Cellular Imaging & Histology Core. The confocal microscope is equipped with four solid-state lasers line (405, 488, 567, and 637nm) and the appropriate filter cubes. The acquired images were analyzed using ImageJ and QuPath, version: 0.4.3 (University of Edinburg) for colocalization of OT+ and AVP+ cells with c-fos expression. Immunohistochemistry was also performed for the examination of anterograde tracing, and confirmation of viral expression in the optogenetic stimulation studies.

#### Whole brain activity mapping

C57Bl6 virgin female subjects were each exposed to one of the four temperature conditions for one hour. Within 5-15 minutes after the brain tissue were collected after standard 4% PFA transcardial perfusion followed by an overnight incubation in 4% PFA at 4°C. Samples were washed three times for 1 hour each with PBS and stored in PBS + .02% Sodium Azide at 4°C for storage and shipment. The whole brain tissue samples were sent to Transluscence Biosystems for whole brain activity analysis. The sample were cleared using iDISCO method. Then, probed using immunoglobin against markers for neural activity (immediate early genes, c-fos and Npas4). Images were acquired using light sheet imaging on ZEISS Lightsheet Z.1 and Lightsheet 7 microscopes using our Mesoscale imaging System™. This was followed by machine learning-based image analysis with data registration using the Allen Brain Atlas as reference. The data was analyzed based on fold change in cold and warm temperature conditions relative to the control temperature conditions (RT).

#### Anterograde tracing

To examine the brain regions that receive inputs from PVN-AVP cells, stereotaxic viral injections were performed in virgin female AVP-IRES-Cre mice. Mice were weighed, then anaesthetized with 1.5–2.5% isoflurane (adjusted based on reflexes and breathing rate during surgery), placed into a stereotaxic apparatus (NeuroStar, Germany). Local anesthetic, Bupivacaine, was injected subcutaneously prior to incision to access the skull. Unilateral craniotomies performed over left PVN (coordinates: anteroposterior (AP): −0.70 mm from Bregma; mediolateral (ML): +0.18 mm from the midline, dorsoventral (DV): −4.7 mm). Cre-inducible, transsynaptic herpes simplex viruses 1 strain, H129ΔTK-TT (National Institute of Health (NIH)- The Center for Neuroanatomy with Neurotropic Viruses (CNNV)) was injected at a depth of 4.7 mm with a 5 µl Hamilton syringe and a 33 gauge needle [84]. The virus was injected into PVN at 0.1 µl min^−1^ for a final injection volume of 0.5µl. The craniotomy was sealed with a silicone elastomer (World Precision Instruments), the skin sutured and antibiotic ointment was applied. Post surgery, analgesia, buprenorphine extended-release injectable suspension was given to subjects at the appropriate dose. Brain tissue samples were collected from subjects after transcardial perfusion 48 hours after surgery to examine anterograde tracing viral expression. Tissue was processed for immunohistochemistry and images acquired with fluorescence microscopy.

#### Optogenetic stimulations: surgery (injections and implantation) and behavior protocol

For optogenetically manipulating PVN-AVP→CEA projections, stereotaxic viral infections and fiber implantations were performed in virgin female AVP-IRES-Cre mice. Subjects were weighed, then anaesthetized with 1.5–2.5% isoflurane placed into a stereotaxic apparatus (NeuroStar, Germany). Local anesthetic, Bupivacaine, was injected subcutaneously prior to incision to access the skull. Bilateral craniotomies performed over left PVN (coordinates: AP: −0.70 mm, ML: +0.18 mm, DV: −4.7mm) and right PVN (coordinates: AP: −0.70 mm, ML: −0.4 mm, DV: −4.7 mm). Cre-inducible engineered opsin expressing AAV, AAV9-Ef1a-DIO ChETA-EYFP, (addgene) or AAV pCAG-FLEX-EGFP-WPRE-AAV9 (addgene) as control was injected with a 5 µl Hamilton syringe and a 33 gauge needle [85]. The virus was injected into PVN at 0.1 µl min^−1^ for a final injection volume of 0.5µl. Craniotomy for optic fibers and supporting bone screw were drilled. Bilateral implantation of 5mm optic fibers (Doric Lenses) were inserted above the left CeA (coordinates: AP: −1.70 mm, ML: +2.6 mm, DV:-4.1mm) and above right CEA, (coordinates: AP: −1.70 mm, ML: −2.8 mm, DV:−4.05mm) at a speed of 10µm sec^−1^. The craniotomy was gently sealed with a silicone elastomer (World Precision Instruments). The implanted optic fibers were secured to the skull using dental cement (C&B Metabond). We applied antibiotic ointment and sutured the scalp. Immediately after surgery we administered analgesia, buprenorphine extended-release injectable suspension. Animals were used for experiments after three weeks to allow for viral opsin (ChETA) expression in PVN-AVP axonal terminals.

To perform optogenetic stimulations, a blue light laser (OptoEngine LLC.) was connected with patch cord and sleeves (Neurophotometrics LLC.) to the optical fibers implanted above CEA of the implanted AVP-ires-Cre virgin mice. At the beginning of each behavior assay, we stimulated with 5ms light pulses, delivered at 20 Hz with laser power between 7 to 15 mW lasting for 30 minutes. Then, the virgin was disconnected to continue the behavior assay. The ‘light off’ experiments were conducted with the laser turned off. Each animal was tested under ‘light on’ and ‘light off’ conditions on separate days, and across mice we alternated the order of on and off days. After behavior studies were concluded brain tissue samples were collected from subjects after transcardial perfusion 48 hours after surgery to examine viral expression and implantation accuracy. The brain tissues were processed for immunohistochemistry as described above, and images acquired via fluorescence microscopy.

## ACKNOWLEDGEMENTS

We would like to thank Dr. Tibor Rohacs (Rutgers), Dr. Vanessa Routh and Dr. Troy Roepke for advising on experimental design and data analysis. We thank Dr. Vineet Chitravanshi for access to the cryostat and Dr. Luke Fritzky and the Imaging Core at NJMS for microscopy training and assistance. We would also like to thank Dr. Taiga Abe and John Cunningham (Columbia University) for their help with behavioral analysis that was not included in this manuscript.

This work has been funded by the Whitehall Foundation and the National Institute of Mental Health (R01 MH128688).

## AUTHOR CONTRIBUTIONS

ZA and IC designed the experiments, analyzed data and wrote the manuscript. ZA conducted the experiments. RO had a major contribution to histology data collection. DBG, ZZM, HKH, BS and NDA contributed to behavioral data analysis. JSR analyzed the whole-brain activity data, contributed to the general design of the study, and edited the manuscript.

## SUPPLEMENTARY FIGURES

**Figure S1:**
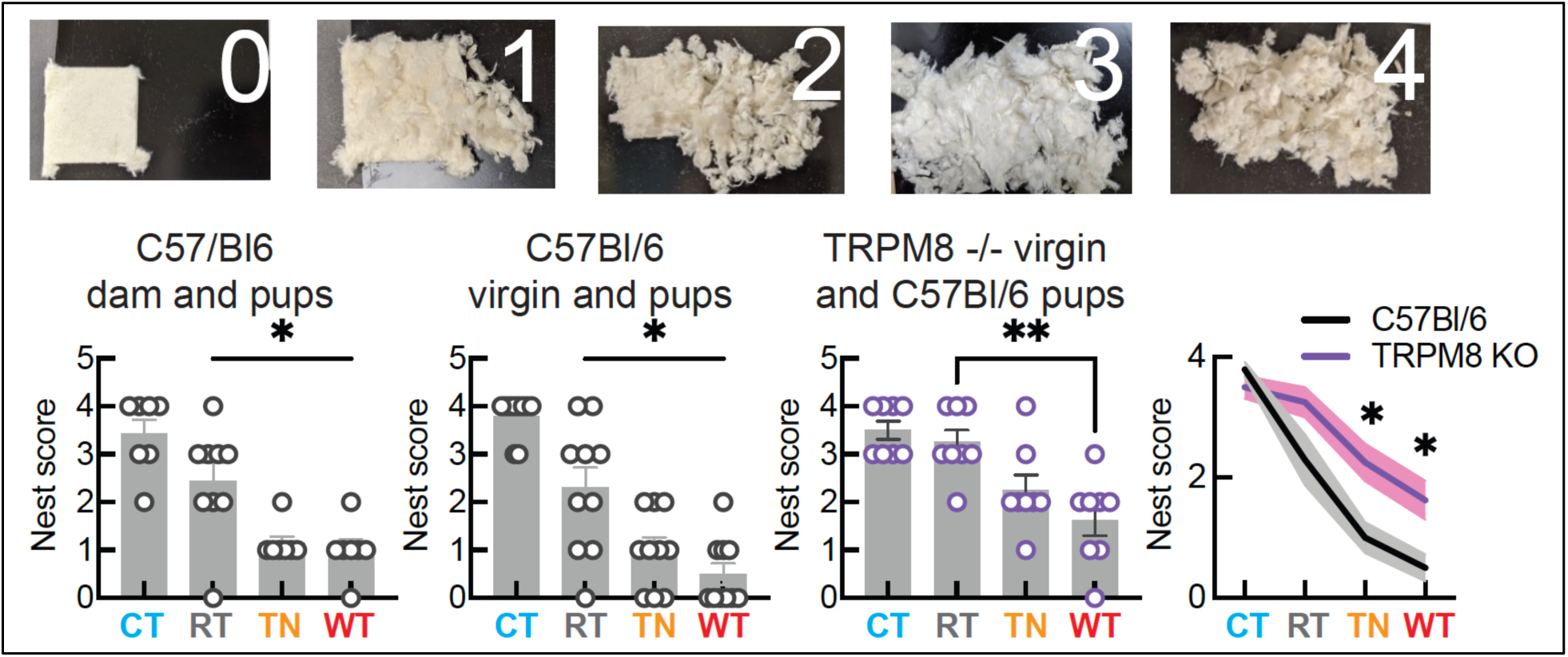
Nest score. Top, example nestles scored with different scores. Bottom, quantification of nest scores. *, p<0.05; **, p<0.01.

**Figure S2:**
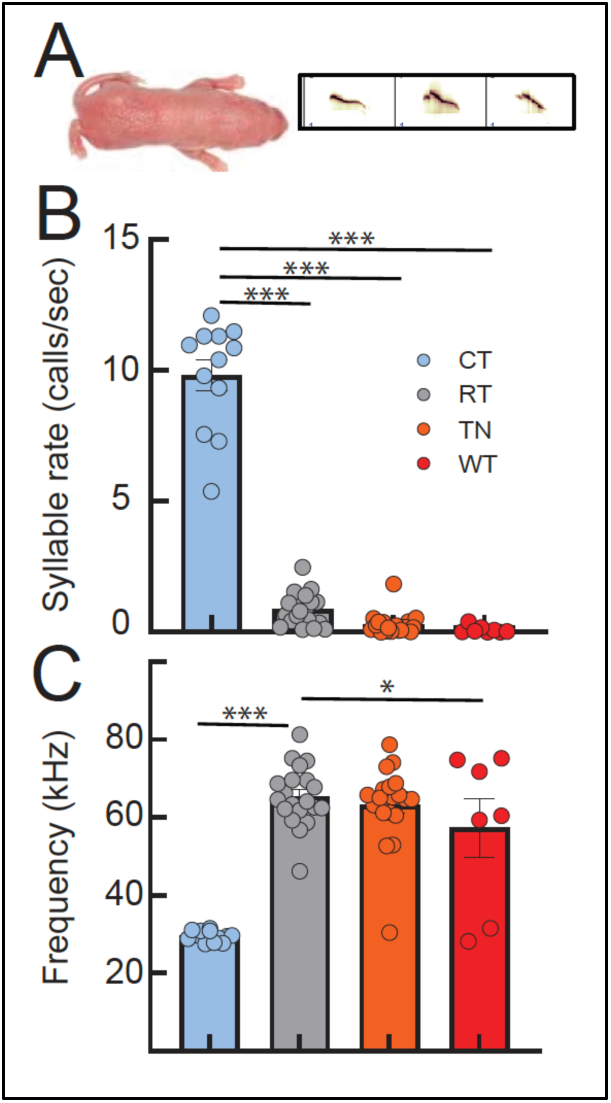
Pup isolation calls in response to changes in ambient temperature. A. Example spectrogram of isolation calls. B. Quantification of syllable rate. C. Frequency of isolation calls. *, p<0.05; ***, p<0.005.

